# The influence of the social environment on larval development and resulting caste in *Bombus impatiens*

**DOI:** 10.1101/2022.09.09.507355

**Authors:** Katherine Barie, Etya Amsalem

## Abstract

The ability of a single genome to express multiple phenotypes is key to understanding social evolution, where individuals with different morphologies perform different tasks. In social insects, phenotypes are typically determined during larval development and depend on the social environment. Here, we used the bumble bee *Bombus impatiens* to examine the social regulation of body size variation and caste by manipulating the identity (queen/worker) and the number of caretakers tending for the brood. Eggs of females and males were kept in cages with (1) a single queen; (2) a single worker; (3) three workers; or (4) ten workers. We measured larval weight, developmental length, and the resulting caste (in females) throughout the brood development of >2000 individuals. We found differences in larval mass when reared by varying number of workers, but not when reared by a single worker compared to a queen. Additionally, in contrast with previous findings in *B. terrestris*, none of the female eggs reared by workers developed into gynes (new queens), indicating that the mechanisms regulating caste determination in *B. impatiens* is not solely dependent on the queen presence as in *B. terrestris*. Adult males were slightly larger than workers and developed for a longer period in the presence of the queen. Overall, we suggest that body size in *B. impatiens* is determined by the number of caretakers and is likely mediated by the amount of provision. The hypothesis that the queen’s presence manipulates female caste was not supported by our data.

## Introduction

Phenotypic plasticity, the ability of one genome to produce multiple variations of behavioral and morphologic forms, is a fascinating phenomenon that is crucial for understanding how societies of social insects have evolved (Corona et al. 2016; West-Eberhard 1989). Diversity of phenotypes, such as females of various sizes and morphologies, is critical for sustaining social organization as they determine the female’s life trajectory as reproductives or helpers.

Social insect societies contain three main castes: queens, workers, and males, while workers can differentiate even further to individuals of different sizes and morphologies (e.g., soldier workers in ants and termites). These differences have been documented in numerous social species (Miura 2005) and form the basis for their social organization. While in many cases, differences between female castes (queens and workers) are limited to body size (Alford 1975; Plowright and Jay 1968; Richards and Packer 1996; Treanore et al. 2020a; Trible and Kronauer 2017; West-Eberhard 1969), female castes in some species exhibit additional morphological differences in the foraging organs or in the spermatheca (Gotoh et al. 2016; Gotoh et al. 2013; Khila and Abouheif 2010). Variation in worker body size and morphology is more common and is often associated with task allocation and reproductive roles. For example, in leaf-cutter ants, the smallest individuals specialize in tending the fungus garden and are more resistant to parasitic fungi (Poulsen et al. 2006; Wilson 1980). Likewise, individuals of the guard caste of the stingless bee, *Tetragonisca angustula*, are generally heavier, have smaller heads and larger legs than forager bees (Smith et al. 2008a). A morphologically-distinct solider caste responsible for nest defense is also common among termites where soldiers have a sclerotized head, enlarged mandibles, a stopper-like shape, or frontal glands which produce defensive secretions (Roisin 2000). The majority of these morphs are determined during early development.

Body size and caste in social insects are influenced by the environment, genetics, or a combination of both (Schwander et al. 2010). In some ant species, such as *Wasmannia auropunctata* and *Vollenhovia emeryi*, caste is genetically determined with the worker caste being produced through sexual reproduction while queens are produced through parthenogenesis (Fournier et al. 2005; Kobayashi et al. 2008; Ohkawara et al. 2006). In Meliponine bees, caste is influenced by a combination of genetic and environmental factors. There, genetic markers in larvae are associated with the queen phenotype but require adequate nutritional input to develop into gynes (future queens) (Hartfelder et al. 2006; Kerr 1950). The same applies to the Florida harvester ant, *Pogonomyrmex badius*, where larval diet differs between castes but patrilineage also influences the resulting caste (Smith et al. 2008b). In many other social species, however, caste and body size are determined solely by environmental factors such as diet, feeding regime, thermoregulation, and colony social conditions (Eyer et al. 2017; Mao et al. 2013; Mutti et al. 2011). In *Vespula maculifrons*, for example, both caste and body size are primarily determined by environmental factors and are not affected by the genetic patriline (Goodisman et al. 2007). Despite extensive study of phonotypic plasticity (Leimar et al. 2012; Libbrecht et al. 2013; Miura 2005; Simpson et al. 2011; Weiner and Toth 2012), our knowledge of the environmental factors determining development, size, and caste is partial and is limited to selected model organisms, primarily ants and some termites, wasps, and bees.

The social regulators of caste and body size are often mediated by the adults, who take care of the immobile young. Adults dictate larval diet, clean and thermoregulate the brood (Jandt et al. 2017; Molet et al. 2017; Yaguchi et al. 2019), and can manipulate not only the larval health but also their life trajectory. Differential feeding in the honey bee, for example, determines whether a female larva develops into a worker or a queen (Eyer et al. 2017; Mao et al. 2013; Mutti et al.

2011), and thermoregulation of brood in social insects has been shown to influence broods’ metabolic rate, growth, caste determination, and health (Howard and Jeanne 2004; Jones and Oldroyd 2006; Kadochová and Frouz 2014; Vogt 1986). The amount of care adults provide depends on the resources available for them, which are driven by the colony size, age and seasonality (Chole et al. 2019; DeGrandi-Hoffman et al. 2018; Hoover et al. 2006; Korb and Hartfelder 2008; Molet et al. 2017; Quezada-Euán et al. 2011; Villalta et al. 2016), but also by their reproductive interests, a theory known as “parental manipulation” in which the caretaker limits the provisioning of the brood to generate smaller or submissive individuals (Alexander 1974b). Parental manipulation by the queen was demonstrated in *Polistes fuscatus* wasps where queens influence larval physiology and caste with vibrational signals in combination with nutritional input (Jeanne and Suryanarayanan 2011; Mignini and Lorenzi 2015; Suryanarayanan et al. 2011). Queens provide wasp larvae fewer vibrational signals, resulting in larvae with a lower likelihood of developing into gynes (Suryanarayanan et al. 2011). Similar works in *Apis mellifera* (Le Conte and Hefetz 2008), *Bombus terrestris* (Cnaani et al. 1997) and *Solenopsis invicta* (Fletcher and Blum 1981) showed that the presence of the queen inhibits the differentiation of larvae to gynes, presumably using pheromones. In some species of termites, the queen is the only individual in the colony able to provide the nutrients larvae require in order to develop as sexuals (Korb and Hartfelder 2008; Yaguchi et al. 2019). Overall, the care received by adults is a strong regulator of larval development and caste in social insects.

Bumble bees are an excellent system for examining the social regulation of body size and caste, because both factors are influenced by the social environment (Cnaani et al. 2000b). However, despite some progress in the field (see below) the mechanisms underlying caste determination are not fully understood. Bumble bee colonies undergo several transitions during the colony life cycle that may result in young receiving differential care. When a bumble bee queen founds a colony, she is the sole caretaker of the brood until the first workers emerge. Female brood in bumble bees is laid by the queen, whereas male brood can be laid by both the queen and the workers (Amsalem et al. 2015). Male and workers brood develop for 24 days, on average, while queens require 36-37 days, on average, in both *B. impatiens* and *B. terrestris*. Eggs hatch within 5 days and go through 4 larva instars, after which they pupate and emerge (Cnaani et al. 2000b; Cnaani et al. 2000c; Cnaani et al. 2002; Tian and Hines 2018). Castes differ mostly in body size (Goulson 2010; Michener and Michener 1974), with queens being 3-4 times larger than workers, whereas males are slightly larger than workers (Goulson 2010; Michener and Michener 1974; Plowright and Jay 1968). Bumble bee workers’ body mass within the same colony can vary up to tenfold (Couvillon and Dornhaus 2010; Cumber 1949), and size variation is often associated with task specialization (Holland et al. 2021; Jandt et al. 2009).

The mechanisms determining female caste and size in bumble bees are not fully understood and the existing data point to a substantial variation across species and sometimes to conflicting data. In *B. terrestris*, the species that was investigated the most, diet composition of provisions is not different for queens and workers (Pereboom 2000). However, queens are fed more frequently than workers in the later stages of larval development (Ribeiro et al. 1999), although this has not been confirmed in a later study (Pereboom et al. 2003). Furthermore, diploid eggs develop to gynes in the absence of the queen, as long as they are separated from the queen before the critical period for differentiation (approximately 5 days after larvae hatch) (Cnaani et al. 1997; Cnaani et al. 2000c). A similar critical period during early development was found in *B. terricola* (Plowright and Pendrel 1977; Röseler 1970), but a much later critical point for differentiation was found in *B. hypnorum, B. rufocinctus*, and *B. ternarius*. In these species it was further suggested that queen determination depends on food quantity and feeding regimes during development (Plowright and Jay 1977; Plowright and Pendrel 1977; Röseler 1970). Queens and workers in these species have similar growth rates but queens are supplied additional food for longer (Roseler 1976).

The frequency of larval feeding in *B. terrestris* and *B. terricola* influences not only caste, but also worker body mass (Pendrel and Plowright 1981; Pereboom et al. 2003). A study that manipulated the pollen intake of *B. terricola* colonies found that reduced food intake of colonies resulted in smaller-bodied workers (Sutcliffe and Plowright 1988). Furthermore, larvae on the periphery of *B. impatiens* colonies are fed less frequently than larvae towards the center of the colony, also resulting in smaller individuals. Variation is worker size is also affected by the presence of the queen. A previous study in *B. terrestris* showed that brood reared by a queen was significantly smaller than brood reared by a worker (Shpigler et al. 2013). A similar study in *B. impatiens* examined development in brood reared by five workers vs. one queen, and found that the queen produced smaller individuals, although whether the effect is due to the identity of the caretaker or their number is not clear (Costa et al. 2021).

In this study, we examined how the social environment, specifically, the identity/caste and the number of the caretaking females affect larval body mass, duration of development and caste in the bumble bee *B. impatiens*. Phylogenetically, *B. impatiens* is much closer to bumble bee species where caste and body size are determined by diet and later in development than to *B. terrestris* where these factors are influenced solely by the presence of the queen (Cameron et al. 2007), but mechanistic details of caste determination in this species are still lacking. To examine this, we grouped female and male eggs with a queen or a varying number of workers (1, 3 and 10). We anticipated that the identity of the caretaker (i.e., queen or worker) would affect female caste. We also expected the number of the caretakers to positively correlate with body size and brood development. Finally, we hypothesized that a selective effect by the queen on females’ body size and development, but not on males, would indicate a parental manipulation aiming to control the resulting female caste and generate submissive and sterile workers.

## Methods

### Bumble bees rearing

Colonies of *B. impatiens* were obtained from Koppert USA, Inc. (Romulus, MI) and were used as a source for caretakers (workers and queens) and egg laying females producing male and female egg batches. Bumble bees lay eggs in batches, containing up to 10 eggs each, and seal them with wax (Amsalem et al. 2015). In all experiments, groups containing 1 and 3 caretakers were kept in small plastic cages (11 cm diameter × 7 cm tall) whereas groups containing 10 caretakers were kept in larger plastic cages (19 × 16.5 × 14 cm) (i.e., to maintain similar density of worker across treatments). Group sizes were chosen based on a previous study showing that reproductive division of labor in groups of 10 workers is similar to a colony (∼40% of the females activate their ovaries and lay eggs), as opposed to smaller groups containing 3 and 5 individuals, where reproduction is monopolized by a single bee (Amsalem and Hefetz 2011). All bees were kept in a dark environmental chamber, at 28-30° C and 60% relative humidity and provided an unlimited supply of pollen and 60% sugar solution.

### Social condition treatments

To examine how caretaker identity affects brood development, we set up cages (n=97) containing an egg-layer queen or a random-age worker together with a single 4-days-old egg batch of either females or males. To examine how the number of caretakers affect brood development, we set up 141 cages with 1, 3, and 10 random-age workers together with a 4-days-old egg batch of either female or male brood. Cages were sampled at different time points, covering the entire duration of brood development (egg to adult). In each cage, we measured larval weight, duration of development, sex, and caste. Caretaker workers were taken from young colonies which were not producing males or young queens. Caretaker queens were taken from colonies upon emergence. They were mated with unrelated males and treated with CO_2_ to induce transition to egg laying as in (Treanore et al. 2021). They were then placed in individual cells until they confirmed to lay eggs. Female eggs were laid by mated queens while male eggs were laid by unmated queens or workers. Cages producing eggs were checked for eggs every 24 hours and were photographed and tagged to keep track of the date eggs were laid. Egg batches were then gently transferred to a treatment cage (i.e., with caretakers) 4 days after they were laid, thus, before hatching and before the earliest critical period for determination (5-6 days after larvae hatch) found in *B. terrestris* (Cnaani et al. 2000b). Workers may lay eggs in the absence of the queen (Amsalem et al. 2015), thus, new egg batches laid by the caretakers after the onset of the experiment were removed daily. Caretakers and egg batches were unrelated across both experiments, since brood care in *B. impatiens* is not affected by relatedness (Starkey et al. 2019).

### Larval development duration, weight, and caste

Brood (eggs, larvae, pupae) or newly-emerged adults were collected daily between days 6 to 26 after the onset of egg laying. At least 10 individuals were collected for each day across all cages in the treatment groups and overall, 2001 brood and newly-emerged adults were collected in the study. The brood was frozen and removed from their wax cases. The developmental stage of the brood (eggs, larvae, pupae, or newly-emerged adults) was determined, individuals were weighed, and the caste was determined for female individuals based on body mass. Workers and males weigh approximately up to 300 mg, while queens weigh >450 mg and can reach up to 1 g. Usually, there is no overlap in the body mass of workers and queens (Amsalem et al. 2015)

### Statistical analysis

Statistical analyses and data visualizations were performed using JMP Pro 16. The effects of the treatments (1, 3, 10 workers or a queen), the time (days since egg laying), and sex (female, male) on the body mass and duration of development of the brood were compared using ANOVA mixed model. Post-hoc comparisons between treatment groups were conducted using Tukey HSD test. The growth rate of larvae across treatments and sexes was compared using indicator function parameterization. Data are presented as means ± S.E.M. Statistical significance was accepted at α=0.05.

## Results

We monitored the development, body mass and caste of 2001 individual brood in the study and examined how the two sexes are affected by the number and identity of the caretaker. Most larvae hatched within 6 days, a slightly older age than in *B. impatiens* (Cnaani et al. 2002), although this could be explained by the method of counting (i.e., whether the first day eggs are found is considered day 0 or 1). The youngest age of larvae was 6 days (from egg laying) while the oldest was 18 (Table 1). The earliest pupation occurred on day 14 and lasted, to the latest, until day 26. All brood emerged on days 24-26 (males and workers) and none of the developed brood was of queens. The development of larvae and pupae in this study is similar to the one reported previously for males and workers of *B. impatiens* (Cnaani et al. 2002) and *B. terrestris* females (Cnaani et al. 2000c).

**Table 1.**
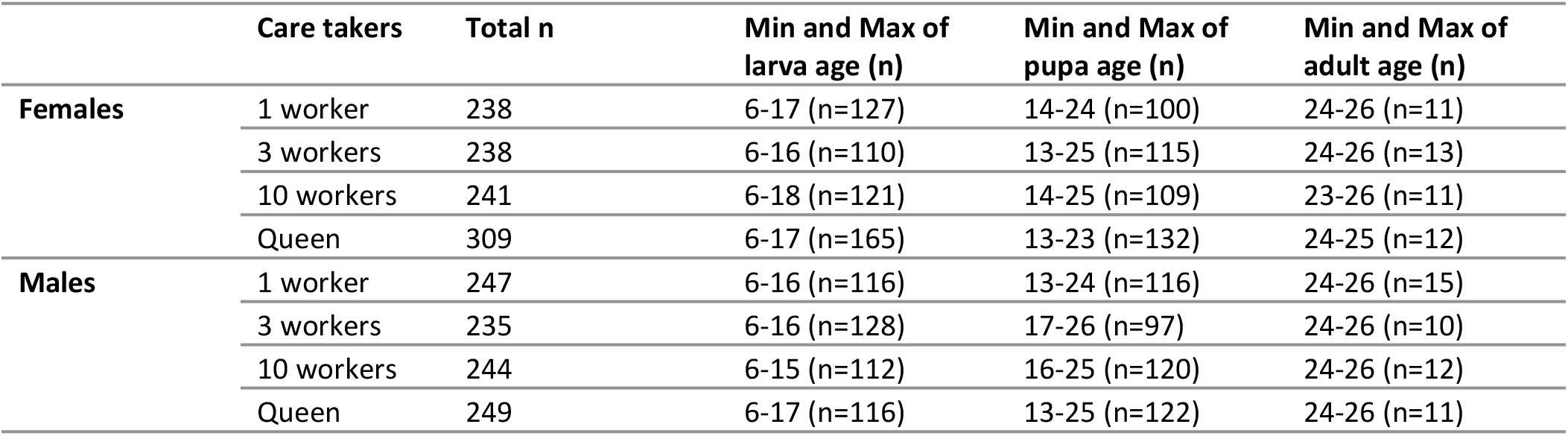
The total sample size in the study split into treatments and sex, and the minimum and maximum values for the larvae and pupae duration of development.

Body mass of female larvae, pupae and adults throughout the development was affected by the time (days after egg laying, Mixed model, Two-way ANOVA f_1,1024_=628.5, p<0.0001) and by the treatment (1, 3, 10 workers or a queen, f_3,1024_=39, p<0.001), however a post-hoc test for the variable treatment showed that the significant impact is attributed to the number of caretakers and not to their identity (post-hoc Tukey HSD p<0.01 for all comparisons except one queen vs. one worker, p=0.08, Figure 1A). A queen and a single worker produced approximately similarly-sized larvae (on average 189±13 and 195±10 mg at the peak of development, n=10 and n=15 for one worker and a queen, respectively, means ± SE), while 3 and 10 worker groups produced much larger larvae (on average 231±6 and 252±17 mg at the peak of development, n=10 and n=10 for 3 and 10 worker groups, respectively, means ± SE). The ‘peak of development’ refers to the timepoint (day) with the highest body mass of larvae.

**Figure 1:**
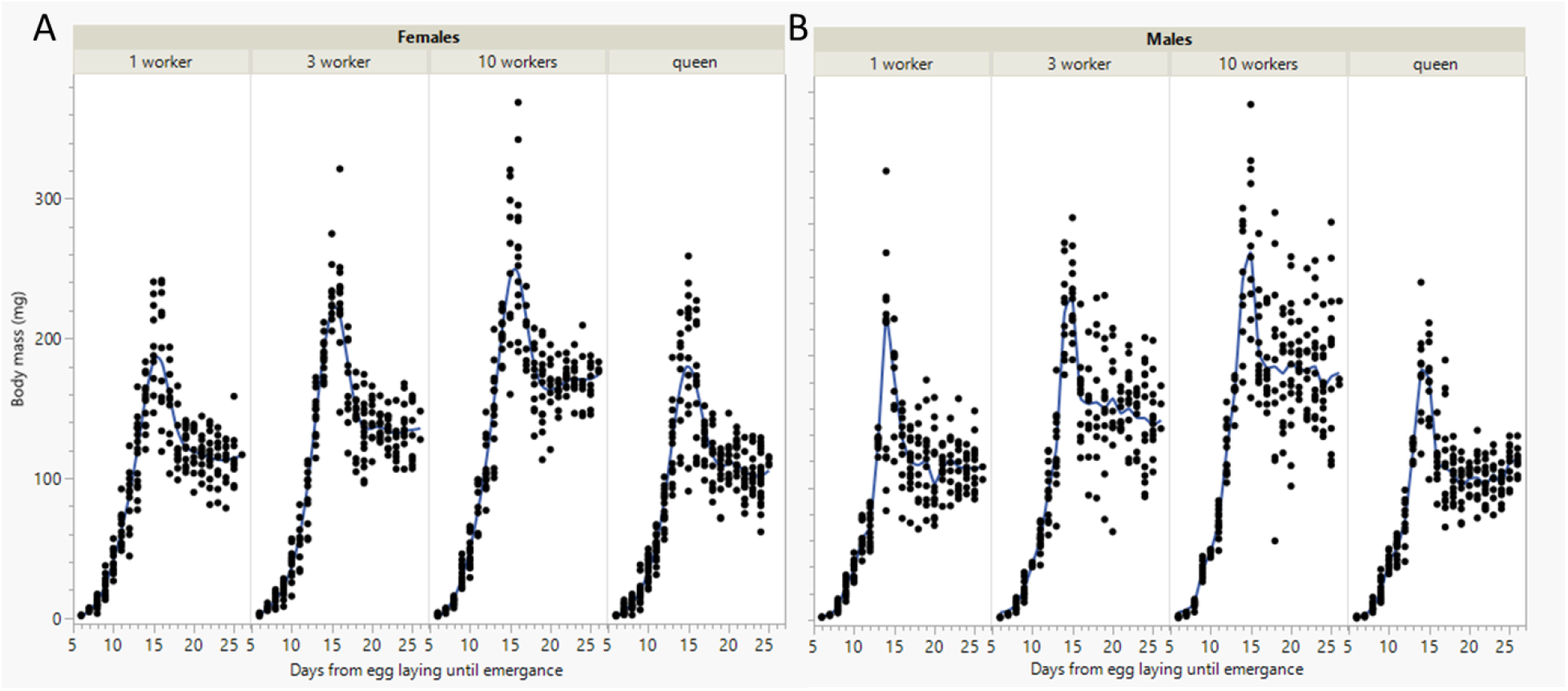
Body mass of female (A) and male (B) larvae and pupae across treatments. Egg batches of either females (laid by a queen) or males (laid by workers) were reared by a single queen or by varying number of workers (1, 3 and 10) and were sampled every day during the entire development period between egg and adult. The sample size per treatment ranges between 235 to 309 individuals (see Table 1 for details).

Body mass of male larvae, pupae and adults was also affected by the time (Mixed model, Two-way ANOVA f_1,973_=614.9, p<0.0001) and the treatment (f_3,973_=59.8, p<0.001), and a post-hoc test showed similar results to females with a significant effect by the number of caretakers but not their identity (post-hoc Tukey HSD p<0.001 for all comparisons except one queen vs. one worker, p=0.64, Figure 1B). Male larvae were overall slightly larger than females (on average 179.4±9, 179.3±10, 243.8±13, and 280±20 mg at the peak of development, n=14, 11, 13, 11 for queen, groups of 1, 3 and 10 workers, respectively, means ± SE), but this difference was not statistically significant (Mixed model with time, sex and treatment as fixed effects, f_1,1999_= 1243.4, p<0.0001 for time; f_3,1999_=97, p<0.0001 for treatment, and f_1,1999_=0.68, p=0.4 for sex, Fig 2).

**Figure 2.**
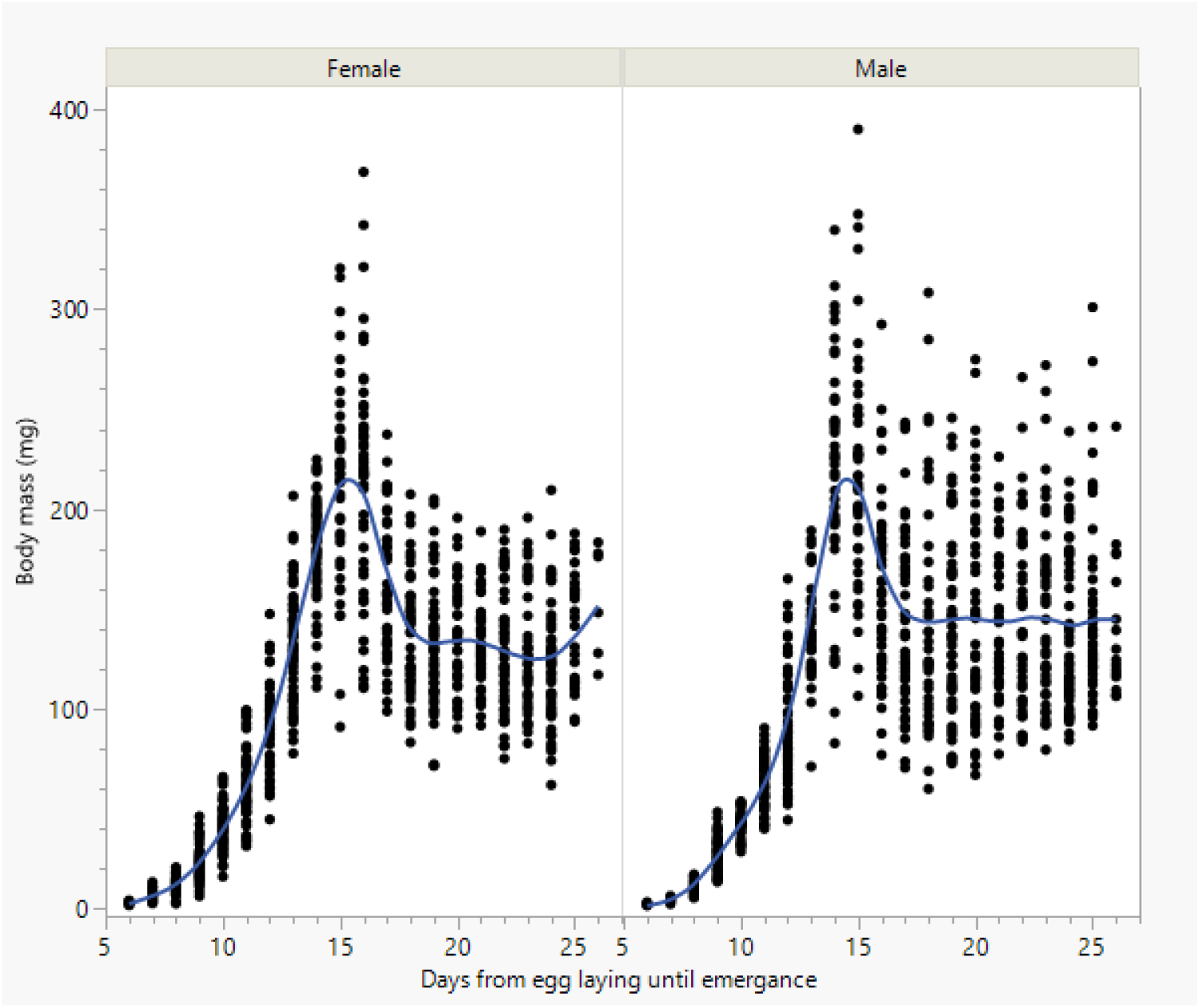
Body mass of female and male larvae and pupae across all treatments. Egg batches of either females (laid by a queen) or males (laid by workers) were reared by a single queen or by varying number of workers (1, 3 and 10) and were sampled every day during the entire development period between egg and adult.

We also separately analyzed the data of female and male larvae to examine differences in their growth rate during the feeding period when reared by a queen and varying numbers of workers. To perform this analysis, we included all the brood that was still larvae at the time of collection (Table 1). This included 523 female and 472 male larvae. The fit of the mass gain curves over time in all treatments and in both sexes was linear (R^2^>0.74). Comparison of slopes in females and male returned similar results to the comparison of the means above, confirming once again that the number of caretakers but not their identity affects body mass gain in larvae (indicator function parameterization for queen vs. one worker: estimate: -3.2, Std Error: 3.66, t-ratio: -0.9, p=0.37; for queen vs. 3 workers: estimate: 9.3, Std Error: 3.8, t-ratio: 2.45, p=0.01; and for queen vs. 10 workers: estimate: 18.9, Std Error: 3.7, t-ratio: 5.1, p<0.001, Fig 3A). Similar analysis of male larvae returned similar results (indicator function parameterization for queen vs. one worker: estimate: 5.8, Std Error: 4.9, t-ratio: 1.18, p=0.23; for queen vs. 3 workers: estimate: 13.1, Std Error: 4.8, t-ratio: 2.7, p=0.007; and for queen vs. 10 workers: estimate: 35.1, Std Error: 4.9, t-ratio: 7.05, p<0.001, Fig 3B).

**Figure 3:**
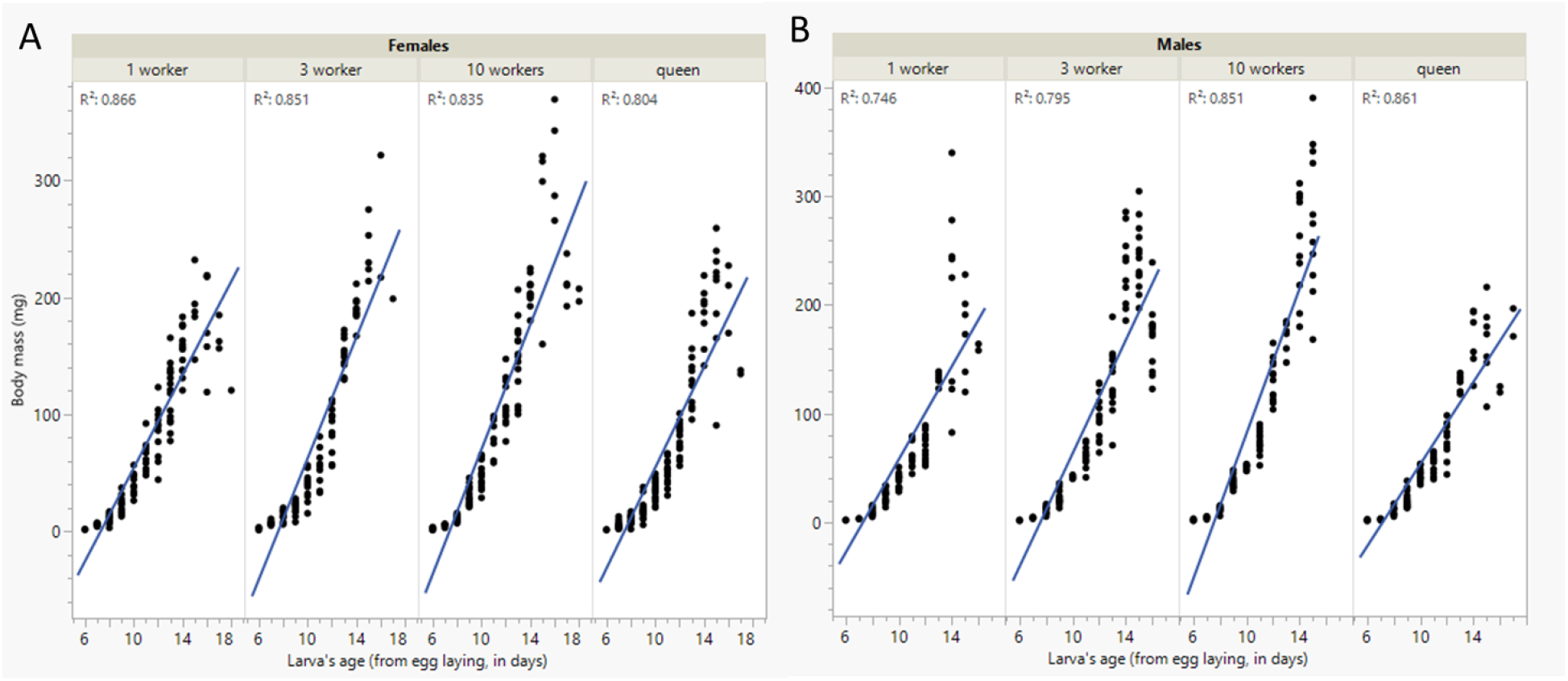
The growth rate of female (A) and male (B) larvae reared by a single queen or by varying number of workers (1, 3 and 10). All larvae were included in the analysis (110-165 individuals per treatment, see Table 1 for details). Larvae hatched on day 6 (after eggs were laid) and pupated on days 13 (the earliest) to 18 (the latest). Most larvae (>95%) pupated on days 15-16.

Finally, we analyzed the effect of treatment and sex on the weight gain and development duration of the brood that emerged throughout the experiment (Fig. 4). This included 47 and 48 female and male adults, respectively (Table 1). Treatment had a significant effect on body mass of females (f_3,45_=27.15, p<0.001 followed by Tukey HSD post-hoc test p=0.6 for one worker vs. a queen and p<0.004 for all the rest) and males (f_3,46_=26.8, p<0.001 followed by Tukey HSD post-hoc test p=0.98 for one worker vs. a queen and p<0.03 for all the rest). Within each treatment, females and males differed in the 10 workers (p=0.01) and the queen (p=0.009) treatments but not in the one worker (p=0.68) or 3 worker (p=0.34) treatments.

**Figure 4.**
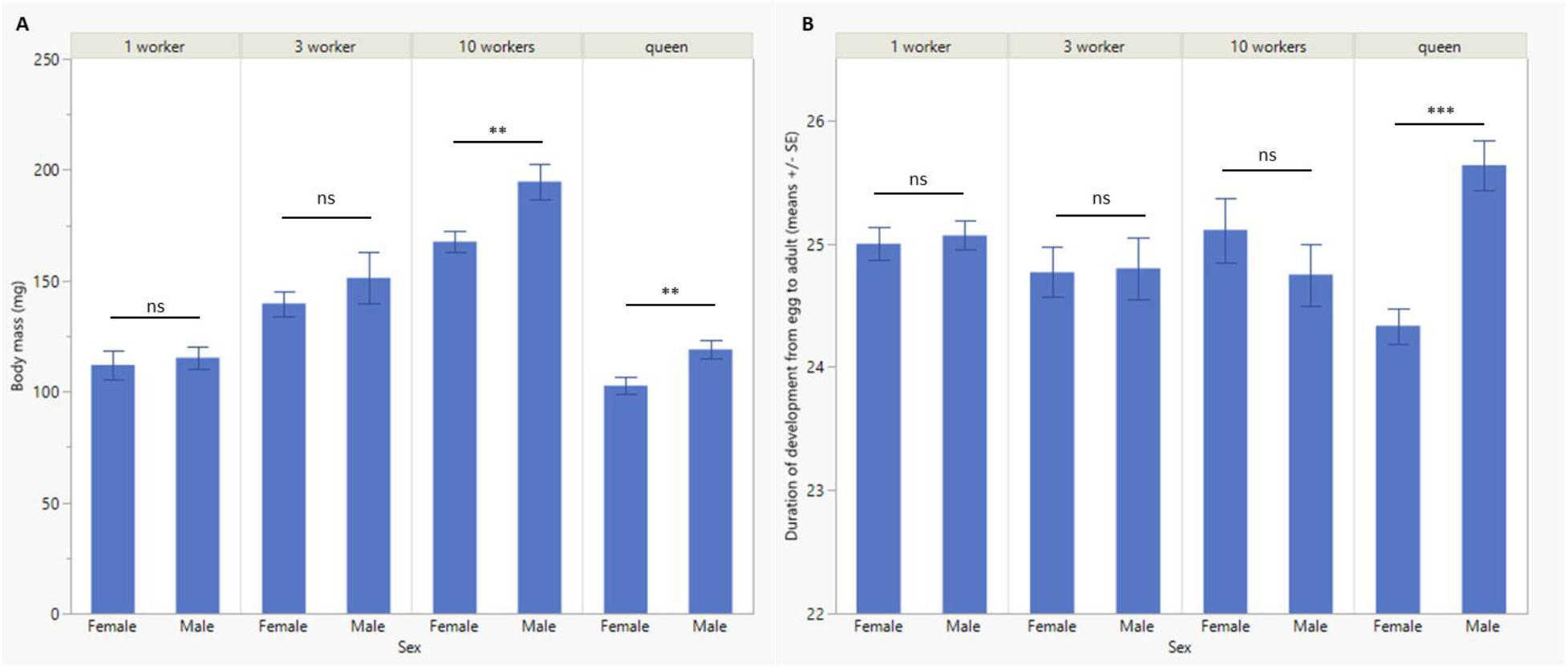
The average body mass (A) and duration of development (B) of all the adults emerging in the study (10-15 individuals per treatment, see Table 1 for details). The body mass was measured upon emergence and duration was counted from the day eggs were laid until the emergence of the adult. Asterisks denote significance at p<0.001 (***) and p<0.01 (**).

The treatment also affected the duration of development in females (f_3,45_=3.37, p=0.02 followed by Tukey HSD post-hoc test p=0.03 for 10 workers vs. a queen and p>0.064 for all the rest) and males (f_3,46_=3.77, p=0.01 followed by Tukey HSD post-hoc test p=0.04 for 3 workers vs. a queen, p=0.01 for 10 workers vs. a queen and p>0.1 for all the rest). Within each treatment, females and males differed in the queen treatment (p<0.001) but not in any of the other treatments (p>0.33).

## Discussion

This study investigated how the social environment affects brood development in *B. impatiens*. We examined the effect of caretaker identity and their number on brood developmental duration, body mass, and resulting caste. Overall, we found that body size is affected by the number of caretakers but not their identity. Additionally, all the female eggs in our study developed into workers, regardless of the identity or the number of the caretakers, indicating that the mechanism determining caste in *B. impatiens* is different than the one in *B. terrestris* and is not solely dependent on the presence of the queen. It is important to note that gynes are typically produced in full-size colonies of *B. impatiens* towards the end of the life cycle, but there is high variability across colonies and the reasons for it are unknown. In a recent study (Santos et al. 2022), we have shown that increasing the amount of the brood in the colony also increases the number of gynes produced by the colony, while decreasing the amount of brood results in colonies specializing in producing males. As far as we know, nobody was able to produce gynes in *B. impatiens* in a setting that is not a full-size colony. These results are fundamentally different from *B. terrestris* where ∼80% of the eggs that were transplanted between queenright and queenless colonies developed into gynes (Cnaani et al. 1997; Cnaani et al. 2000c) and gynes were produced also in small groups of workers (Shpigler et al. 2013). To our knowledge, this is the first time these questions are examined experimentally in *B. impatiens* or in any other species of Bombus that is not *B. terrestris*.

The number of caretakers influenced brood development in a positive way, with an increase in the growth rate and emergence weight as the number of caretakers increased. Similar impacts of the colony size have been observed in other social insect species (Korb and Hartfelder 2008; Molet et al. 2017; Plowright and Jay 1968; Plowright and Pendrel 1977; Ramalho et al. 1998). These effects are likely mediated via an increase in feeding frequency, attributed to the number of caretakers (Pereboom et al. 2003; Ribeiro et al. 1999). Interestingly, the size differences between female and male brood were minor and apparent only in the adults reared by the queen and not throughout development (Fig. 4). In contrast, *B. terrestris* males were reported to be larger and fed more frequently than workers (Ribeiro et al. 1999; Shpigler et al. 2013). However, an analysis of *Bombus* species found a large overlap in the size range of males and workers, and although males are generally larger in *B. terrestris*, this is not the case in *B. impatiens* (Del Castillo and Fairbairn 2012).

We found no effect of caretaker identity on the developmental time, growth rate, and weight at emergence, with the brood reared by a single worker being similar in all parameters to the brood reared by a queen. Based on the parental manipulation hypothesis, it is predicted that the queen would manipulate brood development to produce smaller workers in order to reduce competition with her offspring (Alexander 1974a). Accordingly, we expected that queen-reared brood would develop quicker and be smaller than worker-reared brood as was found in *B. terrestris* (Shpigler et al. 2013). However, since the queen has no competition with males, we expected this effect to be specific to female eggs. Conversely, we found that a queen-reared brood was not significantly smaller nor developed faster than a worker-reared brood. The difference in body size of brood reared by workers and queens in *B. terrestris* could be the result of worker rearing gynes in the absence of the queen. However, decreased developmental time in individuals reared by the queen was observed even when no gynes were produced, suggesting that in *B. terrestris*, but not in *B. impatiens*, the queen is able to manipulate worker size. In line with these findings, a previous study in *B. impatiens* also found no differences in the developmental time of female brood reared by a queen versus brood reared by five workers (Costa et al. 2021). However, this study did find that a single queen produced smaller individuals compared to the worker social condition, but this finding may be confounded by the number of caretaker (a single queen versus 5 workers) (Costa et al. 2021). Potentially, both queens and workers in *B. impatiens* may preferentially produce smaller individuals when there are a few caretakers because it may be less costly and require less provisioning to produce smaller individuals (Cnaani and Hefetz 2001; Couvillon and Dornhaus 2010).

Regardless of caretaker identity or number, no gynes were produced in any of the social condition treatments. The absence of a queen has been shown to incite rearing of gynes in several social insects such as the ant species *Aphaenogaster senilis* and *Atta sexdens* and honey bees (Boulay et al. 2009; Tarpy et al. 2000; Winston 1991). Additionally, in *B. terrestris*, close to 80% of the female brood that was reared in the absence of the queen and that was transplanted before the critical period for differentiation, developed into gynes or intercastes in both full-size colonies (Cnaani et al. 2000a), and also in small groups of 10 workers (Shpigler et al. 2013), and increased colony size affected the production of new gynes in several studies (Bloch 1999; Pereboom et al. 2005). However, in this study we found an increase in brood body mass (within the range of a worker) with the increase in the number of caretakers but did not observe gyne production even in the ten-caretaker condition, meaning that the queen is not the sole factor affecting caste determination and gyne production.

That being said, a group of ten workers may not be sufficient for inducing gyne production in *B. impatiens* and an increase in the number of nurses combined with the absence of the queen may have resulted in gyne production. A previous study in several bumble bee species has suggested a certain worker/larva ratio needs to be achieved for gyne production (Plowright and Jay 1968). This ratio, however, may vary across bumble bee species. In some ant species it has been reported that the number of workers constrained reproductive decisions, and the production of gynes was lower in smaller than in larger groups. In the instances of low worker numbers, rearing gynes may require an overhead that small worker groups cannot afford (Ruel et al. 2012).

Since we observed no difference in brood reared by one worker and one queen, our data do not indicate that there is an active manipulation of the queen on worker body mass or duration of development. The data further show that no gynes were produced in any of the worker cages, questioning whether the queen’s presence is as important for gyne production in *B. impatiens* as it is in *B. terrestris*. Data in *B. impatiens* support a much simpler model in which worker size (and maybe also caste) are dependent on the number of caretakers and consequently, the amounts of resources. Size variation in workers can be explained along the same lines with the production of relatively smaller individuals either early in the season, when fewer caretakers are available, or very late in the season, when resources are no longer available (Treanore et al. 2020b). *B. impatiens* may have a similar caste determination mechanism to that seen in *Bombus hypnorum, Bombus rufocinctus*, and *Bombus ternarius*, to which they are more closely related (Cameron et al. 2007) and where caste determination depends on food quantity and feeding regimes during development (Plowright and Jay 1977; Plowright and Pendrel 1977; Roseler, 1970; Roseler and Roseler 1974). However, the detailed mechanisms determining gyne production in this species are yet to be explored.

## Acknowledgments

We thank the MS committee members of KB (DRs Heather Hines, Christina Grozinger and Rudolf Schilder) for reading and providing feedback on previous drafts of the manuscript.

## Declarations

### Ethics approval

A license or certificate was not required through Pennsylvania State University for experiments involving bees. Nevertheless, we conducted all experiments in accordance with international animal care and use ethical standards.

### Availability of data and material

The datasets used and/or analyzed during the current study are available from the corresponding author on reasonable request.

